# An interaction network of RNA-binding proteins involved in *Drosophila* oogenesis

**DOI:** 10.1101/2020.01.08.899146

**Authors:** Prashali Bansal, Johannes Madlung, Kristina Schaaf, Boris Macek, Fulvia Bono

## Abstract

During *Drosophila* oogenesis, the localization and translational regulation of maternal transcripts relies on RNA-binding proteins (RBPs). Many of these RBPs localize several mRNAs and may have additional direct interaction partners to regulate their functions. Using immunoprecipitation from whole *Drosophila* ovaries coupled to mass spectrometry, we examined protein-protein associations of 6 GFP-tagged RBPs expressed at physiological levels. Analysis of the interaction network and further validation in human cells allowed us to identify 26 previously unknown associations, besides recovering several well characterized interactions. We identified interactions between RBPs and several splicing factors, providing links between nuclear and cytoplasmic events of mRNA regulation. Additionally, components of the translational and RNA decay machineries were selectively co-purified with some baits, suggesting a mechanism for how RBPs may regulate maternal transcripts. Given the evolutionary conservation of the studied RBPs, the interaction network presented here provides the foundation for future functional and structural studies of mRNA localization across metazoans.

## Introduction

The post-transcriptional regulation of gene expression requires several *trans*-acting factors that regulate the life cycle of an mRNA (1). Many of these factors are RNA-binding proteins (RBPs) that interact with the maturing mRNAs to form functional messenger ribonucleoprotein complexes (mRNPs), interconnecting various steps of RNA metabolism, thereby controlling gene expression (1–3). In *Drosophila* oogenesis, mRNPs are first assembled in the nurse cell nucleus, providing a platform for the formation of larger dynamic assemblies in the cytoplasm, regulating mRNA transport, silencing and localized translation. Several RBPs have been identified and extensively studied in *Drosophila* development. Some of the well characterized and evolutionary conserved examples include the dsRNA-binding protein Staufen (Stau) (4–11), the DEAD-box helicases Vasa (Vas) (12–20) and eIF4AIII (21–25), the CCHC-type zinc finger protein Nanos (Nos) (26–30) and the heterogeneous nuclear ribonucleoproteins (hnRNPs) Hrp48 and Glorund (Glo) (31–33). Binding of these proteins is essential for the proper expression of four key maternal transcripts: *bicoid (bcd), oskar (osk), gurken (grk)* and *nanos (nos)*, critical to define the future embryonic axes.

During *Drosophila* oogenesis, the anterior-posterior axis is established through the localization of *bcd* to the anterior pole and localization of *osk* and *nos* to the posterior pole of the oocyte. Accumulation of *grk* at the antero-dorsal corner determines the dorso-ventral axis of the embryo. The posterior targeting of *osk* requires several RBPs including Stau, eIF4AIII, Hrp48, Glo and Vas (4, 25, 34–38). Once localized, translation of *osk* initiates the assembly of the pole plasm by anchoring Vas to the posterior of the oocyte, a critical step in the formation of germ cells (39–41). This also results in posterior localization and activation of *nos*, essential for the embryonic abdominal patterning (42). Remarkably, many components of the *osk* mRNP regulate multiple transcripts. For example, Stau is also essential for the anterior accumulation of *bcd* mRNA in the eggs (43, 44). Hrp48, Glo and Vas regulate the localization and translational regulation of both *osk* and *grk* transcripts (35–38, 40, 45–47). Glo also represses non-localized *nos* in the oocytes (48), while Vas promotes *nos* translation in the embryos (49).

In addition to their functions in establishing oocyte polarity, these RBPs have various other roles during *Drosophila* oogenesis. For example, Hrp48 and Glo are required in nurse cells for the regulation of chromosome organization (38, 47). They have also been implicated as regulators of alternative splicing, consistent with the role of their mammalian homologs (38, 50–52). During early oogenesis, both Vas and Nos are involved in the maintenance of germline stem cells, in oocyte differentiation and other aspects of oocyte development (26, 35, 53–56). In embryos, Nos functions in germline development (53, 57–61) and further promotes the inclusion of germline cells in the developing ovary (53, 62). In addition to oogenic processes, Nos and Stau are also involved in the development of the *Drosophila* nervous system (63, 64).

Many RBPs in *Drosophila* oogenesis have overlapping functions that are likely differentially regulated. Little is known about this regulation and it may involve several as yet unidentified mRNP components. To comprehensively identify RBP interactors, we carried out a systematic *in vivo* purification screen of GFP-tagged RBPs coupled with mass spectrometry. We employed both labeled and label-free MS methods and identified several proteins significantly enriched with the purified RBPs. The interactomes of the individual RBPs were largely independent with some overlap. Our screen identified several previously unknown interactions, many of which we validated *in vitro*. This work presents an extended interaction network of RBPs in *Drosophila*, offering a new reference point for future functional and structural studies of mRNA localization.

## Experimental procedures

### Cloning and DNA constructs

For cloning purposes, total RNA was extracted from wild-type ovaries using TRI-Reagent (Sigma), according to the manufacturer’s instructions. RNA was reverse transcribed using Moloney Murine Leukemia Virus (M-MuLV) reverse transcriptase (Thermo Fisher Scientific), in the presence of oligo (dT)_15_ primers. To express proteins in human HEK293 cells, genes of interest were amplified from *Drosophila* cDNA, or in some cases from the Drosophila Genomics Resource Center (DGRC) clones, using the standard PCR conditions. Accession numbers of all the genes cloned are provided in Table S1. Fragments were cloned into mammalian expression vectors based on pEGFP-C1 (Clontech), bearing an N-terminal EGFP tag or modified to contain either HA or HA-Flag tags (provided by Elisa Izaurralde, MPI Tübingen). Full-length cDNAs were cloned, except for protein Nucampholin (Ncm), where a sequence encoding amino acids 359-664 was amplified. The boundaries were designed based on the MIF4G domain of the human ortholog CWC22, which has been shown to bind eIF4AIII (65). To serve as a control, either MBP or EGFP alone was used.

### *Drosophila* stocks

All flies were kept at room temperature on standard *Drosophila* medium. Oregon R flies were used as wildtype. Fosmid lines expressing GFP-tagged proteins were purchased from the Vienna *Drosophila* Resource Center (VDRC 318283, 318719, 318195, 318157, 318898, 318766), as previously published (66). To generate the control fly line expressing the tag only, the tag sequence was cloned in a modified pUAST-attB vector (67) (without UAS sites or SV40 poly(A) signal), downstream of a moderately expressing *exu* promoter using *Kpn*I and *Bam*HI sites. The purified vector was injected into embryos from a recombinant stock with a genotype y[1] M{vas-int.Dm} ZH-2A w[*]; PBac{y[+]-attP-3B}VK00033 (BDSC 24871). Transgenic flies were identified in the F1 generation by the presence of red eyes (dsRed) and a stable fly line was established.

### Cell culture and co-IP from HEK cells

Human HEK293 cells were grown in standard Dulbecco’s Modified Eagle Medium, supplemented with 10% heat-inactivated Fetal Bovine Serum, Glutamine and Penicillin-Streptomycin solution, at 37 °C in the presence of 5% CO2.

For co-immunoprecipitations (co-IP), transfections were carried out in six-well plates with Lipofectamine 3000 (Invitrogen) according to the manufacturer’s recommendations. Typically, 5 *μ*g of DNA was transfected in each well and the ratio of two plasmids was adjusted based on their expression levels. If required, a third empty plasmid with HA tag was supplemented, to reach a total amount of 5 *μ*g. Cells were collected 2 days after transfection and washed with PBS prior to lysis. Cells were lysed for 15 min on ice in buffer (50 mM Tris-HCl pH 7.5 at 4 °C, 100 mM NaCl, 250 mM Sucrose, 0.1% NP-40, 1 mM DTT) supplemented with protease inhibitors (cOmplete^™^, EDTA-free Protease inhibitor cocktail, Roche). For efficient lysis, cells were mechanically sheared by passing them through a needle (Sterican 21G 7/8” Ø 0.8X22mm) several times. Cell lysates were cleared at 16,000*g* for 15 min at 4 °C and supernatants were supplemented with 5 μl/ml of RNase A/T1 (Thermo Fisher Scientific) for 30 min at 4 °C. After clearing the lysate again at 16,000*g* for 15 min, 12-20 μl of GFP-TRAP MA beads (Chromotek) were added to the supernatant and the mixtures were rotated for 1 hour at 4 °C. For Flag pull-downs, 1.8 μg of monoclonal anti-Flag (Sigma #F1804) was added to the supernatant, post RNase treatment. After 1 hour at 4 °C in rotation, 20 μl of GammaBind Plus Sepharose beads (GE Healthcare) were added, and the mixtures were rotated for an additional hour at 4 °C. Beads were washed with lysis buffer and proteins were eluted in sample buffer by boiling at 95 °C for 10 min.

### Immunoprecipitation from *Drosophila* ovaries

Ovaries from well-fed flies were dissected in PBS and stored at −80 °C. For immunoprecipitation, frozen ovaries were thawed on ice in lysis buffer (50 mM Tris-HCl pH 7.5 at 4 °C, 100 mM NaCl, 250 mM Sucrose, 0.1% NP-40 and 1 mM DTT) and pooled together in required numbers (see Table S2). Ovaries were mechanically homogenized in lysis buffer (320 μl/ 40 flies), supplemented with protease inhibitors (cOmplete™, EDTA-free Protease inhibitor cocktail, Roche), using tissue homogenizer with a glass pestle. Lysates were cleared by centrifugation at 21,000*g* for 20 min at 4 °C and 5 μl/ml of RNase A/T1 (Thermo Fisher Scientific) was added to the supernatants. After incubation at 4 °C for 30 min, lysates were cleared again and 30-60 μl of GFP-TRAP MA beads (Chromotek) were added. The mixtures were incubated for 1 hour at 4 °C in rotation. Beads were washed with lysis buffer and proteins were eluted as described above.

### Western blotting and detection

Eluates were separated on 10% polyacrylamide gels and transferred to a nitrocellulose membrane. Membranes were blocked in PBS containing 5% milk powder and 0.1% Tween-20. HA-tagged, HA-Flag-tagged and GFP-tagged proteins were detected using HRP-conjugated monoclonal anti-HA (1:5000, BioLegend #901501) or polyclonal anti-GFP antibodies (1:2000, Thermo Fisher Scientific #A11122) respectively. Blots were developed with ECL (GE healthcare) reagents, as recommended by the manufacturer, and imaged using an Amersham Imager 600 (GE healthcare).

### Experimental design and statistical rationale

For analysis of proteins interacting with each tagged RBP, both label-free and dimethyl labeling MS experiments were performed and raw data was processed by MaxQuant software as described below. Proteome data comprised a total of 21 raw files (3 biological replicates from each sample) for label-free MS and 2 raw files (2 biological replicates from each sample) for dimethyl labeling MS. Tag alone was used as a negative control for both analyses.

### Mass spectrometry measurements

For proteome measurements, eluates were separated on a NuPAGE Bis-Tris precast 4-12% gradient gel (Invitrogen). Samples were run approximately 2 cm into the gel and bands were visualized with a 0.1% Colloidal Coomassie Blue stain (Serva). Proteins were digested in-gel using trypsin. Peptides were desalted and purified on C18 StageTips (68). LC-MS analysis were performed on a nanoLC (Easy-nLC 1200, Thermo Fisher Scientific) coupled to a Q Exactive HF mass spectrometer (Thermo Fisher Scientific) through a nanoelectrospray ion source (Thermo Fisher Scientific), as described previously (69). In brief, peptides were eluted using a segmented gradient of 10%–50% HPLC solvent B (80% ACN in 0.1% formic acid) at a flow rate of 200 nL/min over 46 min. MS data acquisition was conducted in the positive ion mode. The mass spectrometer was operated in a data-dependent mode, switching automatically between one full scan and subsequent MS/MS scans of the 12 most abundant peaks selected with an isolation window of 1.4 m/z (mass/charge ratio). Full-scan MS spectra were acquired in a mass range from 300 to 1650 m/z at a target value of 3 × 10^6^ charges with the maximum injection time of 25 ms and a resolution of 60,000 (defined at m/z 200). The higher-energy collisional dissociation MS/MS spectra were recorded with the maximum injection time of 45 ms at a target value of 1 × 10^5^ and a resolution of 30,000 (defined at m/z 200). The normalized collision energy was set to 27%, and the intensity threshold was kept at 1 × 10^5^. The masses of sequenced precursor ions were dynamically excluded from MS/MS fragmentation for 30 s. Ions with single, unassigned, or six and higher charge states were excluded from fragmentation selection.

For dimethylation labeling, after tryptic in-gel digestion, derived peptides were loaded on C18 StageTips and labeled as described (70). Measurements were done the same way as for the unlabeled.

### Data Processing and analysis

For label-free MS, raw data files were processed using MaxQuant software suite v.1.6.0.1 (71) at default settings. Using the Andromeda search engine (72) integrated in the software, the spectra were searched against UniProt *D. melanogaster* (taxonomy ID 7227) complete proteome database (11/07/2017; 23300 protein entries; https://www.uniprot.org/), a database comprising a sequence of the tag alone and a file containing 245 common contaminants. In the database search, Trypsin was defined as a cleaving enzyme, up to two missed cleavages were allowed. Carbamidomethylation (Cys) was set as fixed modification, whereas oxidation (Met) and acetylation (protein N termini) were set as variable modifications. The mass tolerances for precursor and fragment ions were set to default values of 20ppm and 0.5Da, respectively. MaxLFQ algorithm was activated and the minimum number of peptide ratio count was set to 1 (73). All peptide and protein identifications were filtered using a target-decoy approach with a false discovery rate (FDR) set to 0.01 at peptide and protein level (Supplementary peptides table) (74). A valid protein identification required at least one peptide with a posterior error probability (PEP) value of 0.01 or smaller. To transfer peptide identifications to unidentified or unsequenced peptides between samples, for quantification, matching between runs option was selected, with a match time window of 0.7 min and an alignment time window of 20 min. Matching was performed only between replicates by controlling the fraction numbers. The same parameters were used to process the raw data from the experiments applying dimethyl labeling, except for the following: MaxQuant software suite v. 1.5.2.8 was used; MS spectra were searched against a reference *D. melanogaster* proteome obtained from Uniprot (16/10/2015; 23334 protein entries; https://www.uniprot.org/); dimethylation on peptide N-termini and lysine residues was defined as light (+28.03 Da), intermediate (+32.06 Da), and heavy (+36.08 Da); re-quantification was enabled; no matching between runs was applied; quantitation of labeled peptides required at least two ratio counts (Supplementary peptides table). Experiments were carried out in biological duplicates (Table S2) and labels were swapped to correct for errors in the labeling procedures.

Bioinformatics analysis of the label-free MS data was done using Perseus v. 1.6.5.0 (75) (Supplementary proteins table for label-freeMS; Supplementary statistics table for label-freeMS). Protein identifications were filtered for potential contaminants, proteins identified only by modified peptides and peptides derived from the reversed sequence of the decoy database. Protein intensity values were logarithmized (Log2) and replicates for each bait were grouped together. For identification of protein interactions, data was analyzed in a pairwise fashion i.e. each individual bait group against the control group. Proteins were filtered based on the identification of min 3 valid values in at least one replicate group. As the data followed a normal distribution, the missing values were imputed (using 0.3 standard deviations width reduction and 1.8 standard deviations downshift) enabling statistical analysis. Both-sided Welch’s t-test was used with s_0_ value of 2, to control the artificial within-group variance. For each test, to filter the rows, a requirement of at least 2 valid values in the bait group was set, further controlling the effects of imputation. A 5-10% FDR cut-off (permutation-based; number of randomizations: 250 without preserving groupings) was set to determine significantly enriched proteins. The same pipeline was employed for all the pairwise analyses.

For analyzing dimethyl labeling MS data, the ratio for each sample was normalized to the median of the distribution through MaxQuant (71), to correct for mixing errors (Supplementary proteins table and analysis for dimethylMS). Normalized ratios were Log2-transformed and ratios from duplicates were plotted against each other. Statistically significant differences in abundance were determined by applying an arbitrary ratio threshold of 1 in Log2 scale (twofold).

All the scatter plots and the volcano plots were generated using GraphPad Prism v.7.0.0. For creating networks or subnetworks, Cytoscape v. 3.7.1 (76) was used. To integrate IP-MS data with literature, information from databases like String v. 11.0 (77) and FlyBase (78) were used. From String, only experimental data with medium confidence range was considered. Physical interaction data from FlyBase (78) were extracted for each protein individually. Node size calculation: For dimethyl labeling data, average of the enrichment ratios of duplicates was calculated for each protein. If a protein was found to be associated with more than one bait or identified in both label-free and labeled MS data, the highest fold change value was considered, irrespective of the experiment type. In cases where several nodes were combined into one, the highest value among the respective individual components was considered. Fig. 4 was created using Gephi v. 0.9.2 (79). Modules were detected with an algorithm described in (80), with randomization on, using edge weights and a resolution of 0.5. Force-field-based clustering was performed using the Force Atlas 2 Plugin. Baits were re-positioned manually for clarity.

For GO term analysis, the DAVID (**D**atabase for **A**nnotation, **V**isualization and **I**ntegrated **D**iscovery) v.6.8 functional annotation tool was used (81, 82), which adopts Fisher’s Exact test to measure the gene-enrichment in annotation terms. The following parameters were used: background: *Drosophila* genome; count threshold (minimum number of genes for that term) of 2; maximum ease score (modified Fisher’s Exact P-value) of 0.01. To reduce redundancy in the GO terms, the DAVID output was fed into REVIGO (**Re**duce + **Vi**sualize **G**ene **O**ntology) (83) and p-values were used to select and cluster GO terms with a similarity score of 0.7 (medium). The *Drosophila* database was used to find the GO term sizes. See supplementary table for GO analysis.

## Results

### Tagged proteins recapitulate the endogenous localization patterns

To purify RBP complexes from fly ovaries under native conditions, we used transgenic fly lines generated by recombineering. We used the “Tagged FlyFos TransgeneOme” resource (fTRG) (66) expressing C-terminally-tagged proteins under the regulation of their endogenous promoters. These lines carry a 40 kDa tagging cassette consisting of “2XTY1-sGFP-V5-preTEV-BLRP-3XFLAG” that can be used for both *in vivo* visualization and affinity purification. From the fTRG library, we selected six RBPs for IP-MS, eIF4AIII, Glo, Hrp48, Nos, Stau and Vas. To serve as a control, we generated a transgenic line expressing the tag alone (hereafter referred to as GFP), under the promoter of a moderately expressing gene (*exu*). To ensure that the RBP fusions are functional *in vivo*, we checked their localization patterns at different stages of oogenesis. All the proteins were found to be localized as expected (Fig. S1a, b). Additionally, we also checked their ability to rescue the effects of mutations that cause either lethality or sterility, as summarized in Fig. S1c. Although only 3 out of 6 transgenes assayed were able to fully substitute for the endogenous copy, their localization in the endogenous patterns suggests that their interactions driving localization during oogenesis have been maintained.

### Label-free MS combined with statistical analysis recovers known associations

For IP, we lysed whole ovaries in mild conditions of salt and detergent, and purified the complexes using the GFP-TRAP system (Chromotek) (Fig. 1a). An IP from flies expressing GFP alone was used as a negative control to identify proteins binding nonspecifically to the tag. Since we were interested in identifying RNA-independent protein-protein interactions, the experiments were carried out in the presence of RNases. As the transgenes are regulated by their endogenous promoters, they had varying levels of expression, as observed by Western blotting (Fig. 1b). To compensate for this variability in the IP-MS analysis, we adjusted the number of flies dissected for each transgene (Table S2).

**Fig 1.**
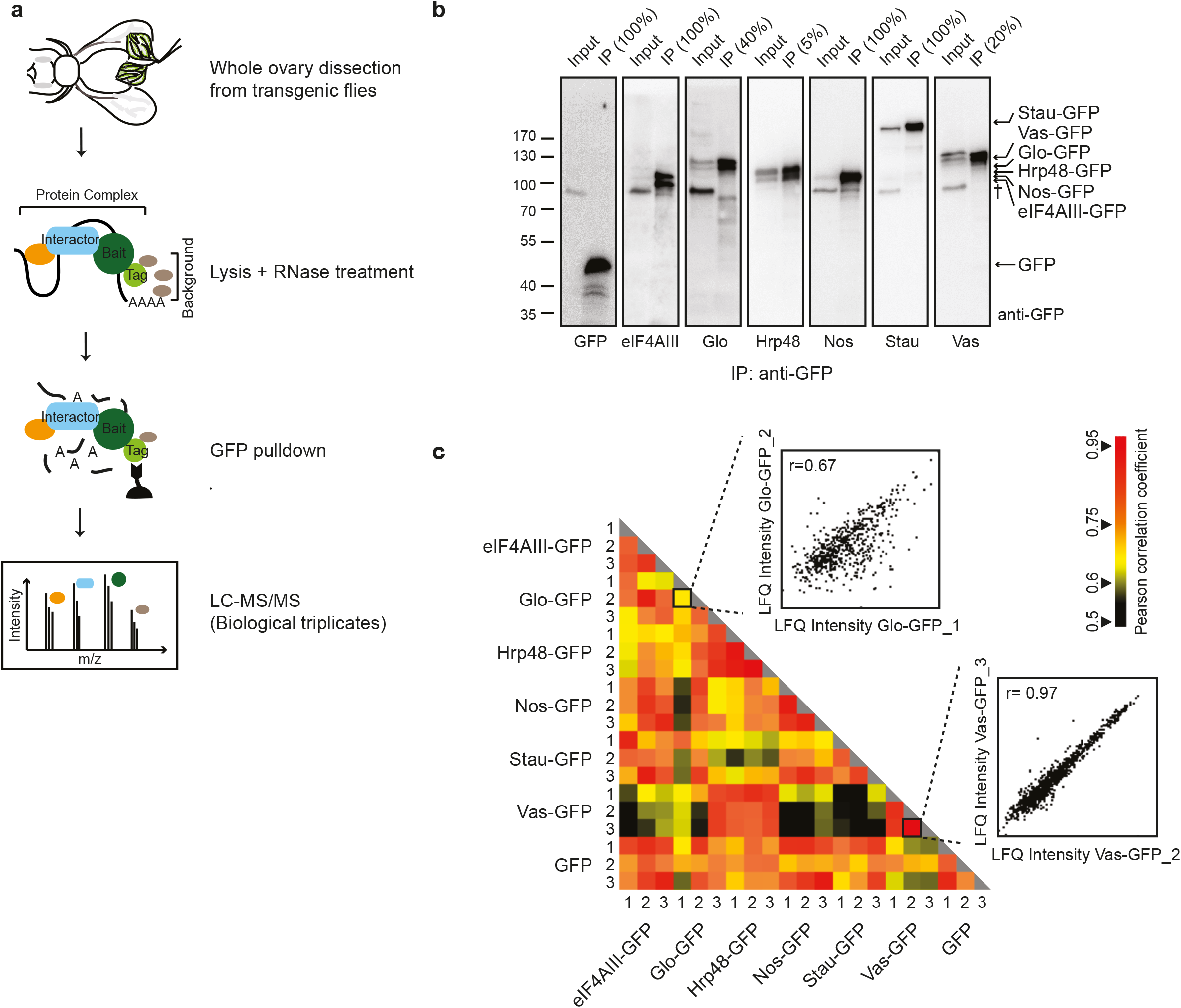
Isolation of bait-associated complexes using IP-MS. (a) Schematic representation of the IP-MS workflow. Whole ovaries were isolated from transgenic flies expressing GFP-tagged proteins (baits) and complexes were purified using an anti-GFP nanobody coupled to magnetic beads. Lysates were treated with RNases prior to immunoprecipitation. IP from a transgenic line expressing GFP served as a control. Eluates were analyzed by LC-MS/MS. (b) Presence of tagged baits in the immunoprecipitates was checked by Western blot, probed with anti-GFP. Inputs and eluates were loaded in different amounts for efficient visualization. Input of the total lysate: 2.1% for GFP, eIF4AIII, Nos; 4.2% for Glo, Vas, Stau and 1.2% for Hrp48. Loading of the respective eluates is highlighted on the top of each lane. † is an unspecific interaction detected by the antibody. (c) Heat map representation of the matrix of Pearson correlation coefficient calculated between LFQ intensities of the complexes, each isolated in triplicate. The zoomed plots show the highest and lowest coefficient values (r) among the replicates.

All the samples were prepared in biological triplicates and the resulting spectra were searched against the *Drosophila melanogaster* proteome database (Fig. 1a, Fig. S2a). For confident identification of proteins and accurate intensity-based Label-Free Quantification (LFQ), we processed the raw data using the MaxLFQ module of the MaxQuant software (71, 73). Additionally, we activated the “matching between runs” algorithm to quantify unidentified or un-sequenced peptides in the samples, by transferring peptide identifications among replicates. The global analysis of the proteomes resulted in the identification of 15,005 peptides mapping to 1878 protein groups, at a FDR of 1% at the peptide and protein level. Of these, 1841 unique protein groups were quantified in at least one of the 21 samples, which account for 87.5% of the total ovary proteome of *Drosophila* (84). The average correlation within replicates ranged from 0.71 (Glo) to 0.92 (Hrp48), suggesting overall good reproducibility of the data (Fig. 1c). In addition, the visualization of LFQ intensities of all the samples as a heat map demonstrates that all the baits were consistently enriched (Fig. S2c). The replicate profiles looked largely similar, with only minor differences. However, the number of proteins quantified with each bait varied highly, as marked by the absence of information in the heat map (Fig. S2c).

For statistical analysis, we considered only those proteins that were quantified in all three replicates of a given sample. To identify significantly enriched proteins, we employed the Welch’s t-test (with the 5-10% FDR cutoff), post-imputation, on each bait-control matrix. The results are presented as volcano plots in Fig. 2a. All the baits were highly enriched and we observed a minimal background, indicating the high specificity of the purifications. For the baits eIF4AIII-, Stau- and Glo-GFP, where fewer proteins were reproducibly quantified, we considered statistical significance up to 10% FDR. The low number of detections for these proteins may be due to the loss of interactions upon RNase treatment.

**Fig 2.**
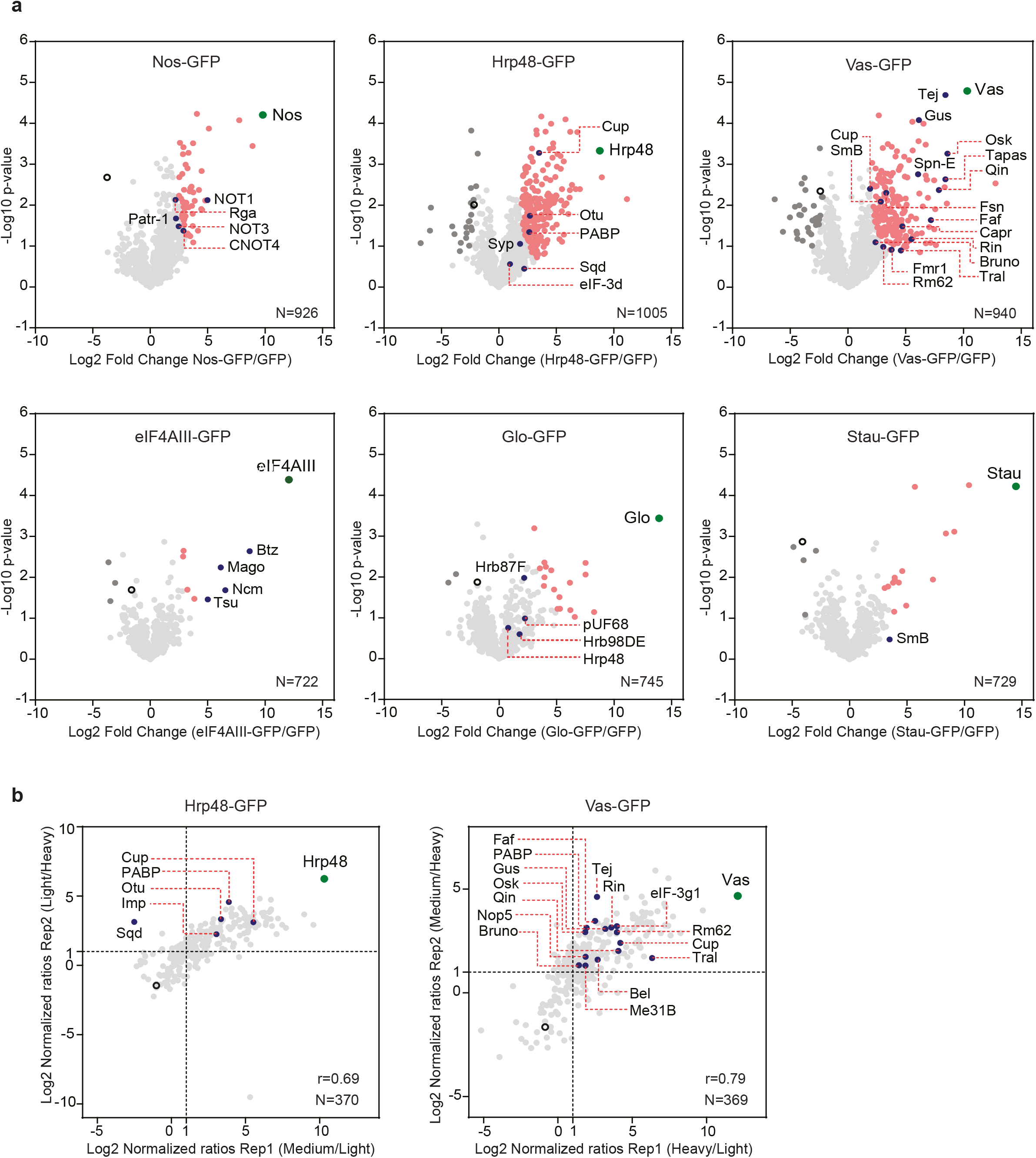
Interactants, including known and new partners, were significantly enriched. (a) Volcano plots of proteins identified to be associated with each bait in the label-free MS analysis, after filtering and data imputation. The significance of enrichment was calculated using the two-tailed Welch’s t-test, with FDR<0.05, s_0=2_ for Nos, Hrp48 and Vas, and FDR<0.1, s_0=2_ for eIF4AIII, Glo and Stau. For each bait-control pair, the resulting differences between the logarithmized means of the two groups “Log2(bait/control)” and the negative logarithmized *p* values were plotted against each other. (b) Scatter plots of the proteins identified to be associated with Hrp48-GFP and Vas-GFP in the dimethyl labeling MS analysis. Normalized ratios (Log2) of both the replicates are plotted against each other. Dotted lines mark the proteins with more than 2-fold change over control, in each replicate. IP from GFP sample served as a control. “N” denotes the number of protein groups plotted and “r” denotes the Pearson correlation coefficient. Each identified protein is represented as a dot in light grey; each bait is highlighted in green; significantly enriched proteins are highlighted in pink; known interactants are highlighted in blue; background binders are highlighted in dark grey; empty circle represents control.

For all the baits, we found several known interactants to be reproducibly enriched over control, mostly with statistical significance (Fig. 2a). For example, we co-purified all the other core components of the Exon Junction Complex (EJC) with eIF4AIII-GFP (25) and the NOT proteins with Nos-GFP (85, 86). We also detected known partners of Vas, involved in both pole plasm assembly in the oocyte (Oskar (Osk); Gustavus (Gus); Fat facets (Faf); F-box synaptic protein (Fsn); Fmr1) and production of germline piRNAs in the nuage (Tejas (Tej); Spindle-E (SpnE); Kumo (Qin); Tapas), with high confidence (87). This indicates that our experimental conditions and analysis pipeline can preserve and identify true interactions. In addition, we also identified proteins that are known to be indirectly associated with Vas, such as the Cullin proteins (88) and Tudor-domain proteins Tudor (Tud) (89) and Krimper (Krimp) (90), suggesting that we not only recovered direct interactants, but whole complexes functional in distinct pathways (Fig. S3). With the Glo-GFP bait, we consistently identified other hnRNPs, including Hrp48 and the splicing factor pUF68, consistent with previous reports (38, 91). However, the enrichment was not significant for these proteins, suggesting weak or transient interactions.

### Quantitative analysis complements the label-free MS data

To detect fold changes in protein abundances with high precision, we used dimethyl labeling MS. This approach is advantageous for *Drosophila*, where metabolic isotopic labeling is still challenging. To confirm the results of our label-free analysis, we carried out dimethyl labeling MS for the Hrp48- and Vas-GFP samples. We chose Hrp48 and Vas as these proteins express well and they are involved in the translational regulation of the same set of transcripts (35, 36, 40, 45–47). We carried out the experiments in duplicates and purified the samples the same way as for label-free MS (Fig. S2b). After in-gel digestion, we labeled the peptides with heavy, medium or light isotopes and inverted the labels in the replicate, to minimize the variability due to the labeling procedures. As before, the raw data were processed with the MaxQuant software (71), providing confident identification of proteins (1% FDR) and normalized protein-abundance ratios. The analysis of the Vas- and Hrp48-associated proteomes resulted in the identification of 4027 peptides, mapping to 615 protein groups. The replicates showed high correlation and the abundance ratios calculated could be well duplicated. We considered as a hit those proteins that we identified with an abundance ratio of >2 in both replicates. Consistent with the label-free analysis, we found several known interactors, most of them reproducibly enriched (Fig. 2b). To check how the two analyses relate to each other, we mapped the proteins identified in labeled MS onto the label-free MS data. As shown in Fig. 3, the proteins that were significantly enriched in the labeled MS followed the same distribution profile and showed up to 47% overlap (for Vas) with those enriched in the label-free MS analysis. Background proteins identified in labeled MS (<2 fold in both replicates) showed a similar profile when graded on the corresponding label-free MS data (Fig. 3). To get a comprehensive view of the proteomes associated with Hrp48 and Vas, we combined the enriched proteins from both analyses.

**Fig 3.**
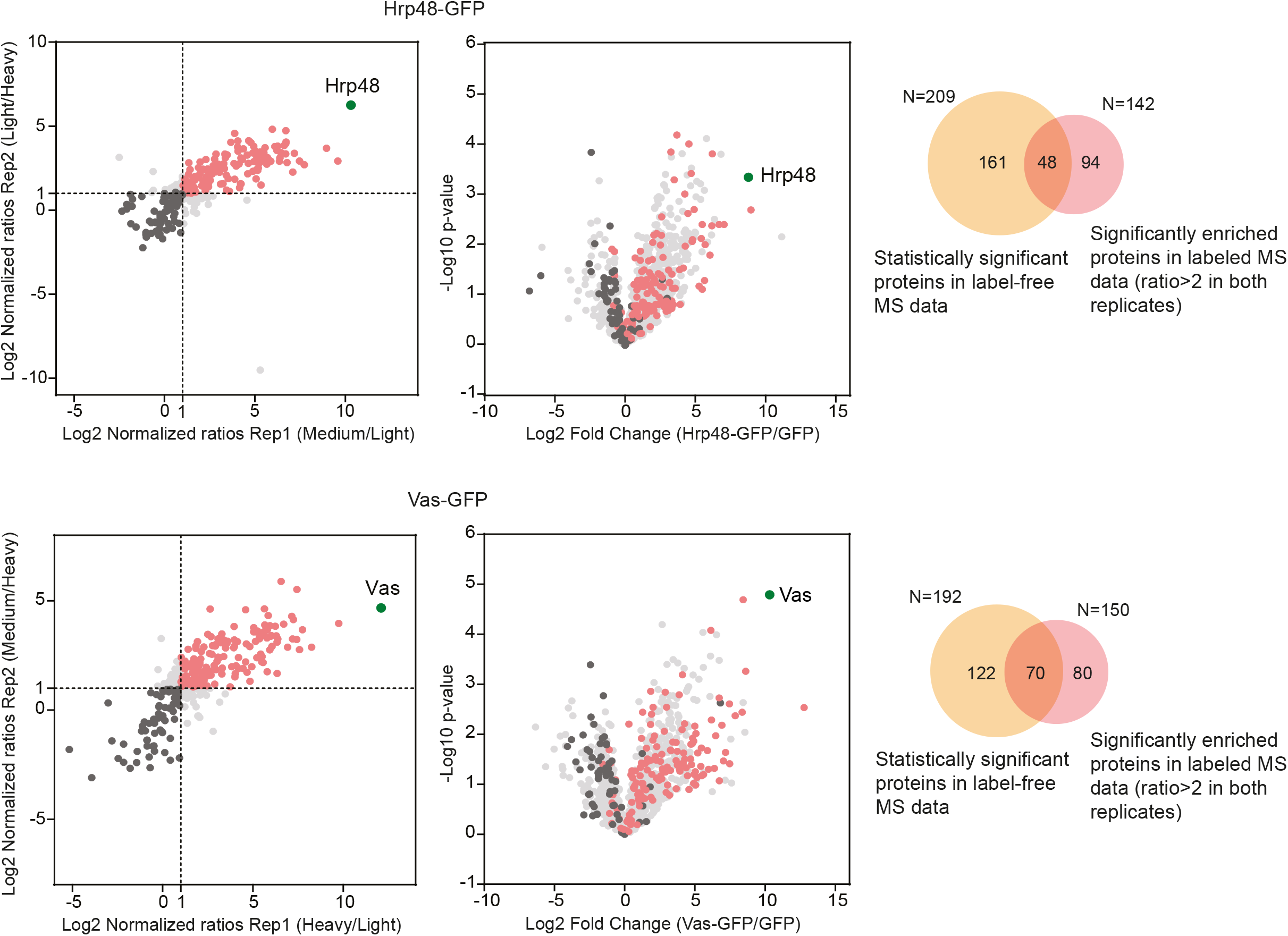
Quantitative label-based MS data support the label-free MS data. Data from the labeled analysis (represented as scatter plots) was compared to the label-free analysis (represented as volcano plots) by overlaying the proteins identified in the respective Hrp48-GFP top and Vas-GFP bottom IP-MS samples. Venn diagrams represent the overlap between the significantly enriched interactants found in the two datasets. “N” denotes the number of proteins analyzed. In the scatter/volcano plots, each identified protein is represented as a dot in light grey; baits are highlighted in green; significantly enriched proteins are highlighted in pink; background binders are highlighted in black.

**Fig 4.**
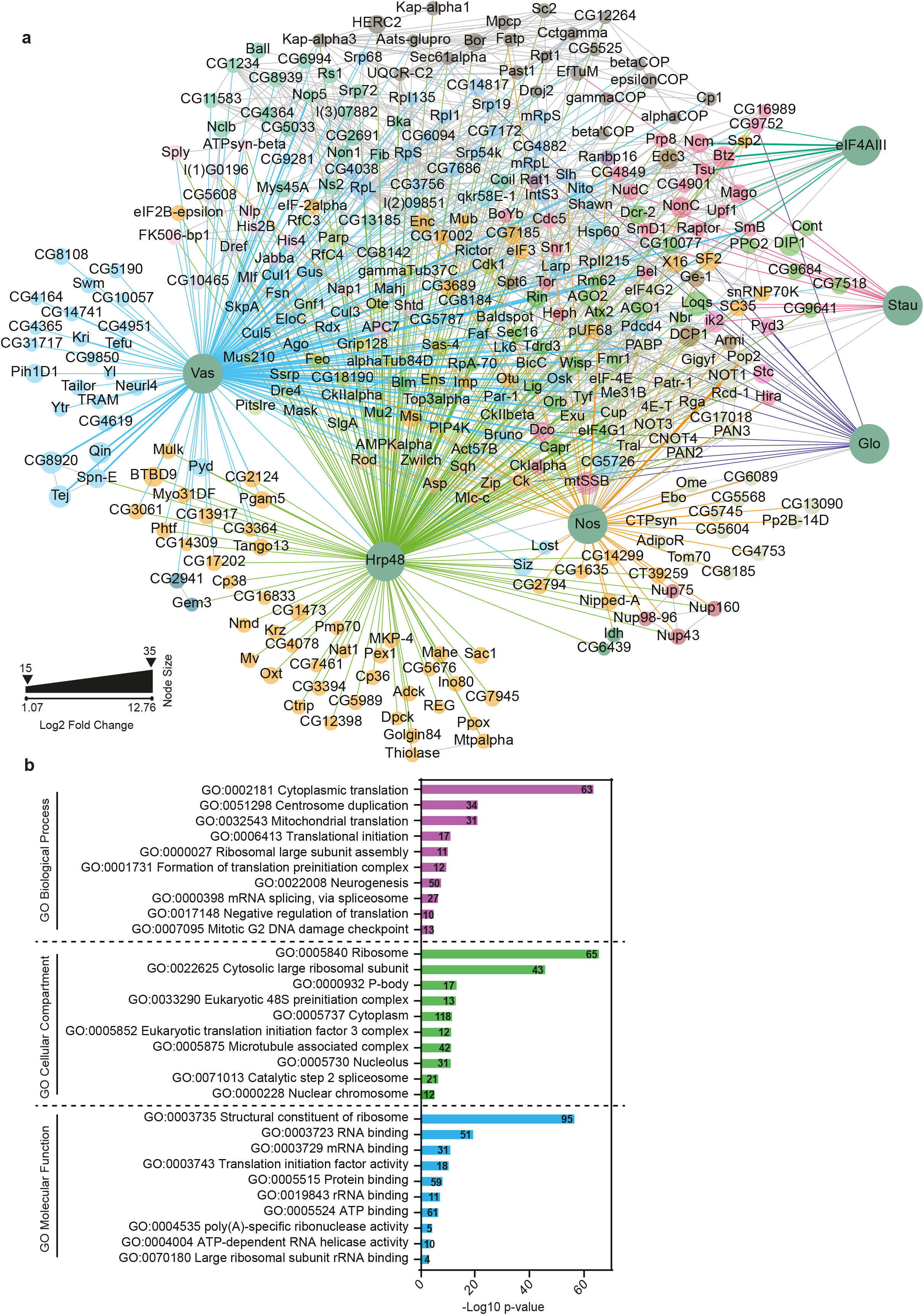
The global interactome reveals a connected network. (a) Interaction network of significantly enriched proteins, identified to be associated with each bait in the MS analysis. For Hrp48 and Vas, proteins from both labeled and label-free analyses were considered. Proteins are represented as nodes while edges represent the interactions. Green nodes represent the baits and the interactants are colored differently, based on their modularity class. The layout is based on a force-field analysis with the baits re-positioned for clarity. Edges representing interactions are colored differently for each bait: Vas in cyan; Hrp48 in green; eIF4AIII in dark green; Glo in blue; Stau in red; Nos in orange and databases in gray. The edges have a unit weight (representing a known interaction). Node size (except for baits) represents the fold change (Log2) over control. For simplicity, respective subunits of the eIF3 complex, cytoplasmic and mitochondrial ribosomal proteins (separately for small and large subunits) were combined into a single node. (b) Proteins were functionally annotated for their roles in biological processes, molecular functions and cellular components using GO term analysis. Only top ten terms in each category, with p<0.01 are shown. The numbers represent the gene count involved in the respective GO term.

### Global analysis of RBP interactomes reveals novel protein interactions

To understand how the proteomes identified with each bait interact with each other, we built a composite network of all statistically significant interactants. While each bait has its distinct proteome, the network is also connected, in agreement with the spatial and temporal distribution of the transcripts they regulate. In particular, we observed a considerable overlap between functionally related proteins such as Hrp48 and Glo (Fig. S4). To gain a systemic understanding of the network, we also added the known protein-protein associations from the String v. 11.0 (77) and FlyBase (78) databases. Next, we carried out a modularity analysis to identify highly connected communities of proteins (Fig. 4a). As expected, many proteins involved in oogenesis, mRNA localization, translational regulation and germ cell formation were selectively enriched with all the baits. Additionally, proteins involved in neurogenesis and splicing were also overrepresented, consistent with the well-studied function of selected RBPs in these processes (Fig. 4b).

Interestingly, we observed ribosomal proteins (components of both large and small subunits) to be significantly enriched with Vas-GFP and Hrp48-GFP, but not with the other RBPs analyzed (Fig. 5, Fig. S4). Previous studies have shown the requirement of Vas in translational activation of *osk, nos* and *grk* mRNAs (35, 36, 40, 49). However, the molecular mechanism by which Vas activates translation is unclear. Studies in *Drosophila* have shown that Vas directly binds the translation initiation factor eIF5B (dIF2) to positively regulate *grk* and *mei-P26* translation and possibly other germline-specific transcripts (92–94). In addition, Vas also interacts genetically with the translation initiation factor eIF4A for efficient germ cell formation (95). However, neither eIF5B nor eIF4A were detected or enriched in our dataset. Instead, other translation initiation factors involved in the formation of the pre-initiation complex, such as eIF2, eIF3 and the cap-binding complex of eIF4E-4G were selectively co-purified (Fig. 5). This is consistent with the recently reported interaction of eIF3 subunits with Vas in the *Drosophila* oocytes (56).

**Fig 5.**
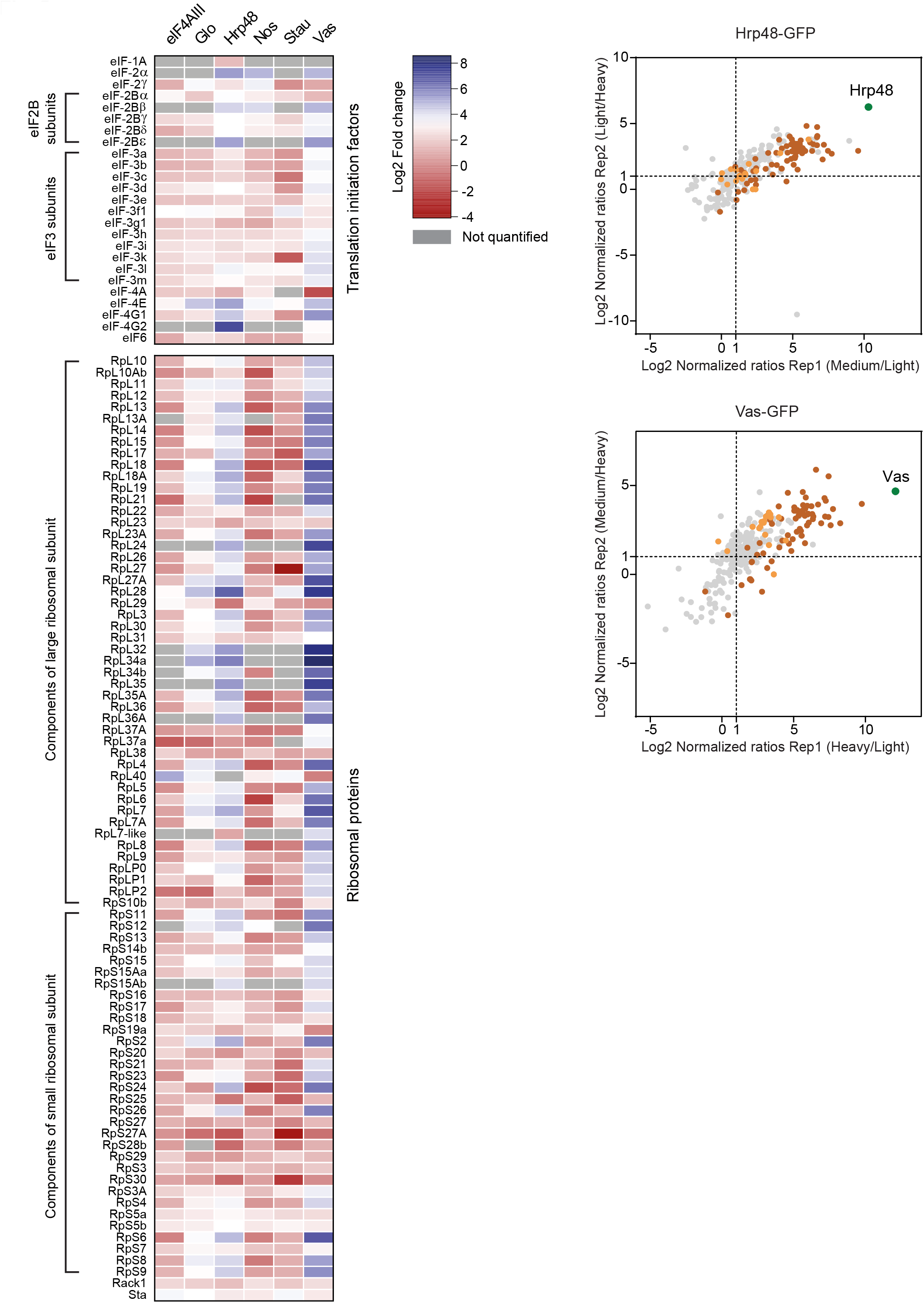
Enrichment of ribosomal proteins and translation initiation factors for Vas-GFP and Hrp48-GFP. On the left, heat map representation of the Log2 fold change of all ribosomal proteins and translation initiation factors co-purified with the six baits, in the label-free MS analysis. Grey rectangles represent empty values. On the right, scatter plots highlighting the distribution of ribosomal proteins and translation initiation factors found associated with Hrp48- and Vas-GFP in the dimethyl labeling analysis. Each identified protein is represented as a dot in light grey; baits are highlighted in green; ribosomal proteins are highlighted in brown; translation initiation factors are highlighted in orange.

Hrp48 is required for the translational repression of *osk* and *grk* mRNAs (45–47). Consistent with this, we copurified with Hrp48-GFP (in both label-free and labeled MS experiments) several P-body-components, associated with the RNA repression/decay machinery, most notably the deadenylase and decapping complexes (Fig. 6). Ribosomal proteins and translation initiation factors were also significantly enriched with the Hrp48-GFP bait, similar to Vas-GFP (Fig. 5). This includes eIF3d, which has recently been reported to interact with Hrp48 to translationally repress the *msl-2* mRNA (96).

**Fig 6.**
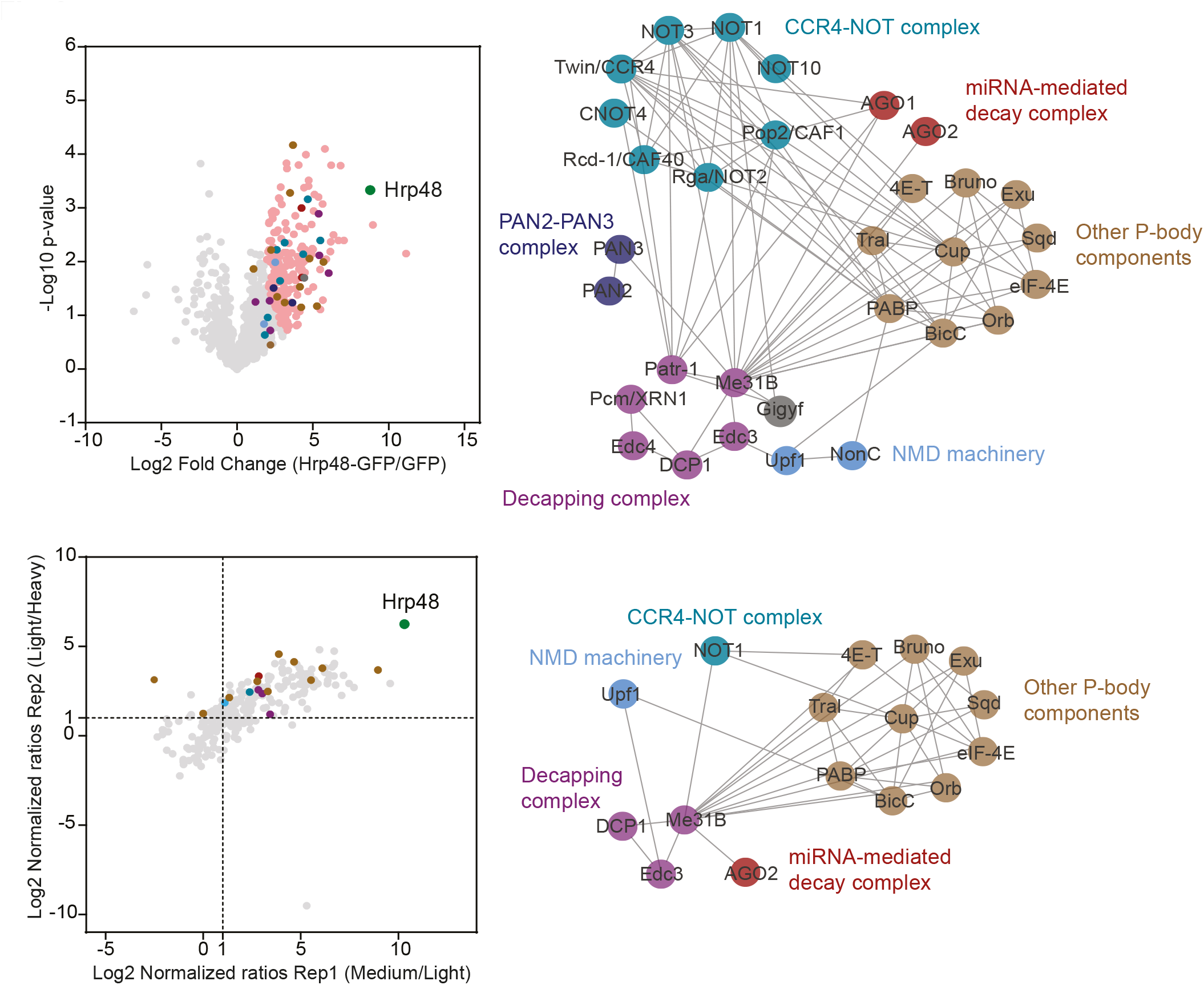
Several components of P-bodies, including the mRNA decay machinery were co-purified with Hrp48-GFP. Volcano plot (top) and scatter plot (bottom) highlighting the distribution of different factors involved in translational repression/mRNA decay associated with Hrp48-GFP in the labeled and label-free MS analyses. Each identified protein is represented as a dot in light grey; the bait is highlighted in green; significantly enriched proteins are highlighted in pink; components of different complexes are highlighted in different colors as indicated in the subnetworks on the right. The subnetworks highlight the interactions between various components of distinct complexes, as previously reported in databases.

### *In vitro* validation of protein-protein interactions

To validate the results of our IP-MS analysis, we co-expressed bait-candidate pairs in cultured mammalian HEK293 cells. This system can be effectively used to study direct interactions of *Drosophila* proteins because it reduces the likelihood of endogenous proteins mediating indirect associations. For validation, we selected significantly enriched candidates identified in the label-free MS analysis. Additionally, we also considered functionally relevant partners, enriched with a >2 times fold change but excluded by statistical filtering. For Hrp48 and Vas, where information from differential labeling was also available, we selected candidates among the interacting proteins identified in both datasets.

Typically, we co-expressed an EGFP (referred to as GFP)-tagged bait with an HA-tagged candidate protein and used the GFP-TRAP system (Chromotek) for IPs. We observed that small proteins (<25kDa) expressed poorly as fusions with HA or HA-Flag. Substitution with the GFP tag improved the expression in all the cases tested. To be able to validate the interactions of such small proteins, we co-expressed an HA-Flag-tagged candidate with a GFP-tagged bait and performed the IP with anti-Flag (Fig. 7f, h). Since Vas does not express well as an HA- or HA-Flag fusion, the small proteins could not be assayed efficiently for interaction with Vas. As negative controls, we used MBP or GFP. We also included known interactions as positive controls, wherever possible. All the tested candidates are indicated in Fig. S5.

**Fig 7.**
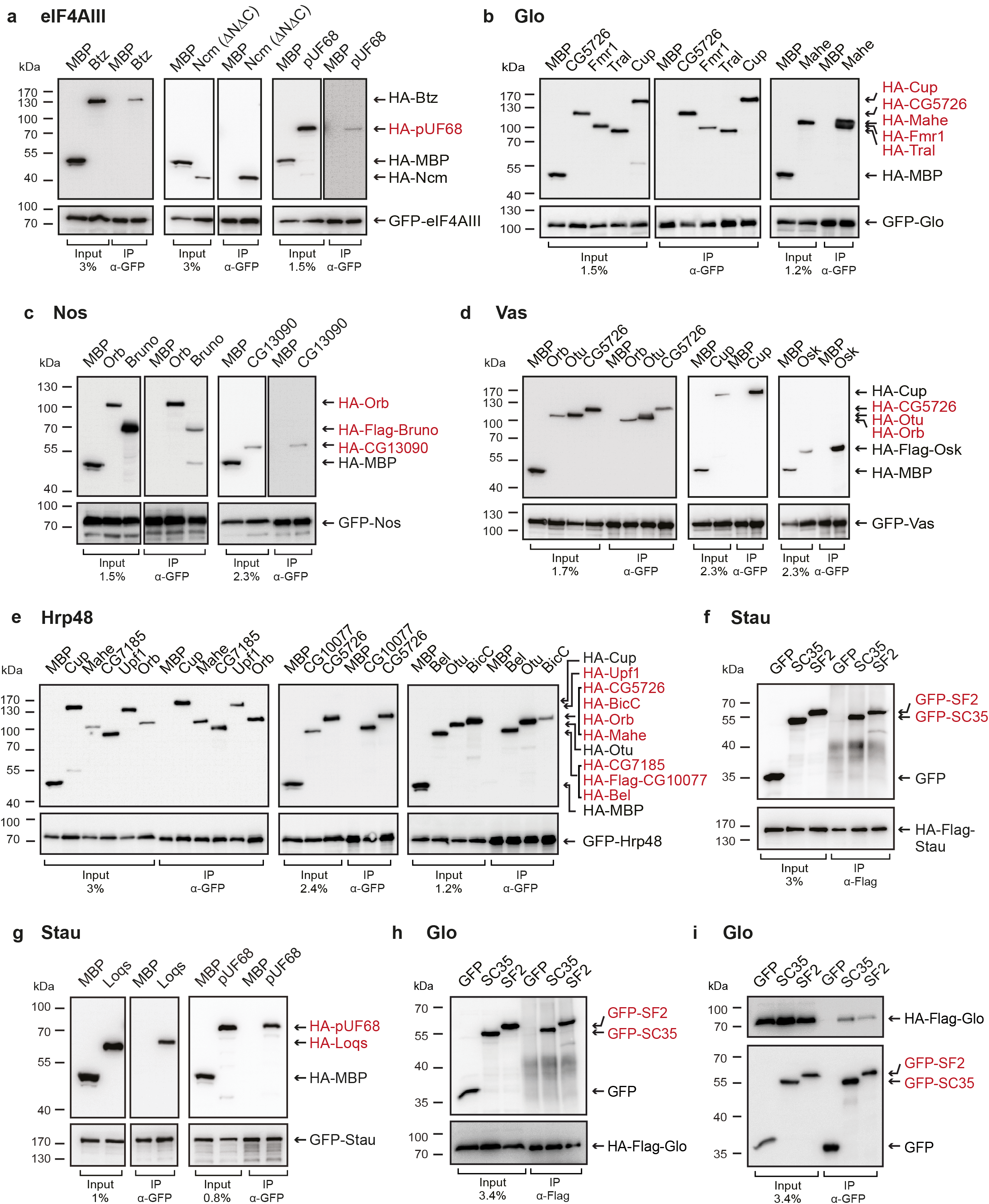
*In vitro* validation of interactions with bait proteins. (a-e, g) Lysates from human HEK293 cells expressing GFP-tagged baits and HA-tagged (or HA-Flag-tagged) candidates were immunoprecipitated using an anti-GFP nanobody coupled to magnetic beads. HA-tagged MBP served as a negative control. Inputs and eluates were analyzed by Western blotting using anti-GFP and anti-HA antibodies. For GFP-tagged proteins, 1-3% of the input and 10% of the eluates were loaded, whereas for HA-tagged (or HA-Flag-tagged) proteins, 1-3% of the input and 90% of the eluates were analyzed. (f, h) HEK293 cells were co-transfected with HA-Flag-tagged baits and GFP-tagged candidates. GFP tag served as a control. Cell lysates were immunoprecipitated with anti-Flag antibody and analyzed by western blotting. For HA-Flag-tagged proteins, 3-4% of the input and 10% of the eluates were loaded, whereas for GFP-tagged proteins, 3-4% of the input and 90% of the eluates were analyzed. (i) Cells were transfected the same way as in panel h, and the lysate was immunoprecipitated with an anti-GFP nanobody coupled to magnetic beads. For GFP-tagged proteins, 3.4% of the input and 10% of the eluates were loaded, whereas for HA-Flag-tagged protein, 3.4% of the input and 90% of the eluates were analyzed. In each panel, cell lysates were treated with RNases before immunoprecipitation. Novel interactions are highlighted in red.

Out of 90 protein-protein interactions assayed, we could confirm 32 interactions (35%), of which 26 were found to be novel (summarized in Fig. S6a). All positive interactions were confirmed at least 3 times, in independent experiments. In addition, we were also able to validate some of the interactions by reciprocal IP, as shown in Fig. 7i. To further confirm that our MS analysis pipeline effectively separated background binders from true interactants, we also tested Nucleophosmin (Nph), which was depleted in all IP-MS datasets. We could not detect interactions of Nph with any of the 4 baits tested *in vitro* (data not shown), in line with the MS data. Additionally, we also tested Sqd that has been reported to interact with Hrp48 in an RNA-dependent manner (47). The negative results further confirm that our experimental conditions effectively disrupted RNA-mediated associations and only protein-protein interactions were confidently identified in our analyses. To visualize the co-IP results, we integrated the validated interactions with the IP-MS data (both labeled and label-free) and information from the literature to create a subnetwork (Fig. S6b). As the majority of the validated interactants are known regulators of maternal mRNAs, this subnetwork highlights a general machinery involved in oocyte development in *Drosophila*.

## Discussion

This study presents a proteome interaction network of six RBPs (eIF4AIII, Hrp48, Glo, Nos, Stau and Vas) required for the localization of maternal mRNAs in *Drosophila*. To construct this network, we purified complexes associated with bait proteins and used the MaxLFQ algorithm (73) for label-free relative protein quantification. The accuracy of this approach is comparable to labeled MS techniques such as SILAC (97). By statistical filtering, we could separate background and specific binders for each bait. Several well characterized interactions were significantly enriched with most baits, indicating the efficacy of our workflow. To complement these data, we also obtained MS data from dimethyl labeling for a subset of the baits. We were able to validate 32 of the interactions assayed, including several novel associations.

In addition to the known regulators of mRNA localization and oocyte patterning, we co-purified nuclear and cytoplasmic complexes involved in different aspects of RNA metabolism. Our results highlight the diverse functions of these complexes in the post-transcriptional regulation of maternal mRNAs. The purification with the Stau-GFP bait of Loquacious (Loqs), a component of the RNAi machinery is one such example. Loqs is a conserved cytoplasmic double-stranded-RNA-binding protein (dsRBP). The protein participates in the biogenesis and processing of small non-coding RNAs that operate within the RNA interference (RNAi) pathway (98). RNAi plays an important role in *Drosophila* germline development and the early phase of *osk* repression (99). *Drosophila Loqs* mutant females are sterile and their ovaries fail to sustain germ-line stem cells (100). We found that both Dicer and Loqs are highly enriched in the Stau IP. We could confirm the Stau-Loqs interaction *in vitro*. This suggests a potential role for Stau in translational repression of *osk* by associating with the RNAi machinery. Although no such evidence has been presented in *Drosophila*, recent reports in other insects (*D. citri* and *L. decemlineata)* (101, 102) and the nematode *C. elegans* (11) have shown a requirement for Stau in RNAi responses. These results suggest a conserved role for Stau in RNAi-mediated gene silencing.

### Nuclear processing is intrinsically linked to cytoplasmic targeting of maternal mRNAs

In addition to their crucial role in pre-mRNA processing, splicing factors also affect the cytoplasmic fates of mRNAs. SmB, a spliceomosomal Sm protein is a known *osk* mRNP component. SmB contributes to germcell specification, at least in part by facilitating *osk* mRNA localization (103) and fails to localize to the posterior of the oocyte in the absence of Vas (104). Several Sm proteins have also been detected to be associated with Vas in the oocytes (56). Consistent with this, we co-purified SmB with Stau, Glo, Hrp48 and Vas. We also purified other splicing regulators with several of our baits, including pUF68 (105) and the SR family proteins SC35 and SF2 (106, 107). Both SC35 and SF2 could be validated for their interaction with Glo and Stau, while pUF68 bound eIF4AIII and Stau *in vitro*. pUF68 was previously shown to interact with Hrp48 and Glo (38) and SF2 co-purifies with the short isoform of Osk (108). These results together with the well-studied role of SR proteins in cytoplasmic regulation of gene expression including mRNA export, decay and translation in mammalian systems (109, 110) suggest that splicing factors are *bona fide* components of the mRNA localization machinery.

### Translational regulation of maternal mRNAs by Hrp48 and Vas

In agreement with the function of Vas in enhancing the translation of maternal mRNAs (35, 36, 40, 49), we co-purified several ribosomal proteins and translation initiation factors with Vas-GFP. Our results suggest that Vas may recruit factors involved in translation initiation. This is also supported by the interaction of Vas with eIF5B and eIF3 subunits, as previously reported (56, 92–94).

In contrast to Vas, Hrp48 is a known translational repressor (45–47). In line with the localization of Hrp48 to P-bodies (111), we co-purified several components of the mRNA decay machinery, including the CCR4-NOT deadenylase complex with Hrp48-GFP. It is possible that these interactions are indirect and mediated by BicaudalC (BicC) and Belle (Bel). These proteins negatively regulate target mRNAs together with the CCR4-NOT complex (112, 113). We demonstrated *in vitro* binding of Hrp48 with both BicC and Bel. This suggests that by recruiting these proteins, Hrp48 may regulate the *nos* and *osk* mRNAs, possibly via CCR4-NOT mediated deadenylation (Fig. 8; ref. (112–115)). The function of Hrp48 in *nos* regulation remains to be investigated (Fig. 8)

**Fig 8.**
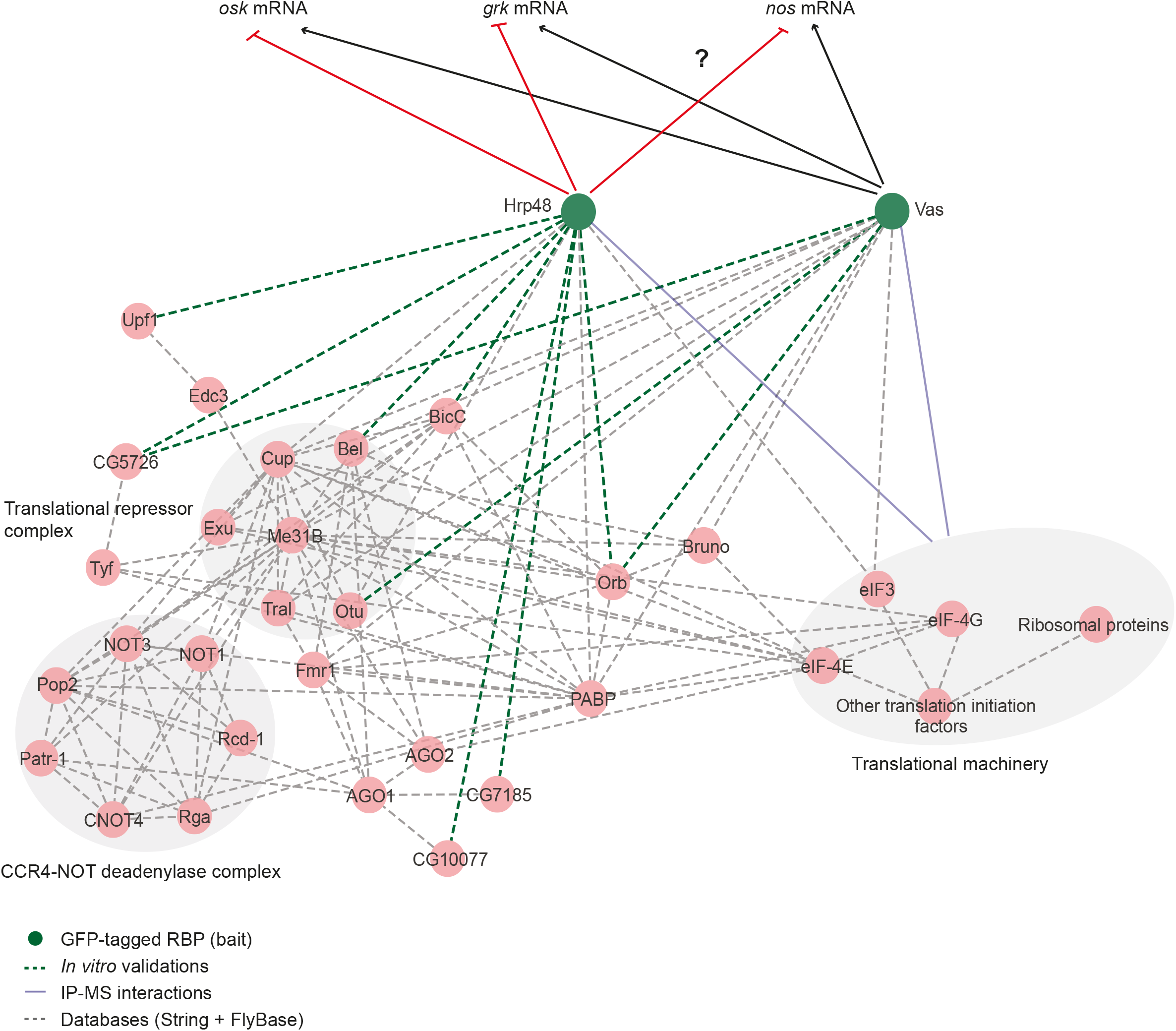
Model of Hrp48-mediated protein associations in the regulation of maternal mRNAs. A subset of protein complexes interacting with Hrp48, possibly involved in the regulation of maternal mRNAs. Many of these proteins were also found to be enriched in Vas-associated complexes. Note that only *in vitro* validated interactions of Hrp48 and Vas are shown here, except for their association with the translation machinery which was identified in the IP-MS data. Proteins are represented as circles and the interactions are represented as edges. Dotted grey lines represent the known interactions from databases; blue lines represent the interaction identified in the IP-MS data; green lines represent the *in vitro* validated novel interactions. Black arrows represent positive regulation while the red arrows represent negative regulation of mRNAs.

However, the parallel enrichment of ribosomal proteins and P-body-components with Hrp48-GFP indicates its bifunctional role in modulating translation. Several lines of evidence from *Drosophila*, including binding of Hrp48 to a derepressor element in the *osk* 5’UTR (45), identification of Hrp48 as a part of a protein complex functioning in translational enhancement of *Hsp83* mRNA (116) and interaction of Hrp48 with CPEB (cytoplasmic polyadenylation element binding) protein Orb (this study) support the dual nature of Hrp48. hnRNP A2, the mammalian homolog of Hrp48, also exhibits the ability to mediate both translational stimulation (117) as well as repression (118), further strengthening the argument.

### Identification of novel genes with a potential role in *Drosophila* oogenesis

Along with several known regulators of maternal mRNAs, we identified the protein products of many previously uncharacterized genes. One such example is CG5726, which encodes for a protein with a MIF4G-like domain. This domain is found in many proteins involved in RNA metabolism including translation initiation factors, NMD factors and nuclear cap-binding proteins (65, 119–122). With no identifiable orthologs in humans, CG5726 protein shows up to 50% sequence identity among Drosophilids. In early embryos of *Drosophila melanogaster*, CG5726 interacts with short Osk (108). In this study, we found CG5726 to be interacting with multiple RBPs: Glo, Hrp48 and Vas, suggesting its possible role in translational regulation.

Consistent with the well-documented role of RNA helicases in oogenesis and fertility (87, 123–126), we identified two putative DEAD-box helicases, CG10077 and Mahe (Maheshvara). While CG10077 is an ortholog of human DDX5, Mahe is an evolutionary conserved regulator of Notch signaling (127). We could recapitulate the interaction of Mahe with both Hrp48 and Glo, and CG10077 with Glo in our co-IP assay. Additionally, we found an interaction of CG13090 (Ubiquitin activating enzyme 4) with Nos. Mutations in this gene have been shown to impair ovarian stem cell functioning (128), which fits well with the role of Nos in maintenance of germline stem cells during oogenesis (53, 55). The functional characterization of the protein interactions uncovered in this study will require further *in vivo* studies.

With certain limitations such as partial functionality of bait proteins or loss of transient or weak interactions, our ability to recover many known associations and validation of several newly identified interactions indicates the general reliability of our data. Although we performed the experiments in a transcript-independent manner, cluster-based analysis of the complexes can be used to identify functional units potentially regulating different mRNAs. Integrating isolated complexes into interaction networks will further enable functional insights into poorly characterized proteins. Given the evolutionary conservation of several RBPs, our study provides a framework to transfer information to other systems, enhancing our understanding of regulation of RBPs and their diverse roles in developmental processes.

## Acknowledgements

We are grateful to Pavel Tomancak and Helena Jambor for their generous gifts of the fTRG fly stocks. We are particularly thankful to Gáspar Jékely for discussion and critical reading of the manuscript. We also thank Uwe Irion for discussions regarding the fly work; Elisa Izaurralde for mammalian cell expression vectors; Desiree Zerbst for providing clones for mammalian cell expression. This project received funding from Max Planck Gesellschaft, the European Research Council (ERC) under the European Union’s Seventh Framework Programme (FP7/2007-2013), ERC grant agreement no. 310957, and the Deutsche Forschungsgemeinschaft (FOR2333, BO3588/2-1 to F Bono).

## Data availability

The mass spectrometry proteomics data have been deposited to the ProteomeXchange Consortium (http://proteomecentral.proteomexchange.org) via the PRIDE partner repository with the dataset identifier PXD016680.

## Abbreviations used

EGFP: enhanced green fluorescent protein
elF: eukaryotic initiation factor
EJC: exon junction complex
FDR: false discovery rate
GO: gene ontology
HA: Hemagglutinin
hnRNP: heterogeneous nuclear ribonucleoproteins
IP: immunoprecipitation
LFQ: label-free quantification
MBP: maltose-binding protein
mRNP: messenger ribonucleoprotein
NMD: nonsense-mediated decay
PFA: paraformaldehyde
piRNA: piwi-interacting RNA
RBP: RNA-binding proteins
SILAC: stable isotope labeling with amino acids
UAS: upstream activating sequence

